# Verification of genetic engineering in yeasts with nanopore whole genome sequencing

**DOI:** 10.1101/2020.05.05.079368

**Authors:** Joseph H. Collins, Kevin W. Keating, Trent R. Jones, Shravani Balaji, Celeste B. Marsan, Marina Çomo, Zachary J. Newlon, Tom Mitchell, Bryan Bartley, Aaron Adler, Nicholas Roehner, Eric M. Young

## Abstract

Yeast genomes can be assembled from sequencing data, but genome integrations and episomal plasmids often fail to be resolved with accuracy, completeness, and contiguity. Resolution of these features is critical for many synthetic biology applications, including strain quality control and identifying engineering in unknown samples. Here, we report an integrated workflow, named Prymetime, that uses sequencing reads from inexpensive NGS platforms, assembly and error correction software, and a list of synthetic biology parts to achieve accurate whole genome sequences of yeasts with engineering annotated. To build the workflow, we first determined which sequencing methods and software packages returned an accurate, complete, and contiguous genome of an engineered *S. cerevisiae* strain with two similar plasmids and an integrated pathway. We then developed a sequence feature annotation step that labels synthetic biology parts from a standard list of yeast engineering sequences or from a custom sequence list. We validated the workflow by sequencing a collection of 15 engineered yeasts built from different parent *S. cerevisiae* and nonconventional yeast strains. We show that each integrated pathway and episomal plasmid can be correctly assembled and annotated, even in strains that have part repeats and multiple similar plasmids. Interestingly, Prymetime was able to identify deletions and unintended integrations that were subsequently confirmed by other methods. Furthermore, the whole genomes are accurate, complete, and contiguous. To illustrate this clearly, we used a publicly available *S. cerevisiae* CEN.PK113 reference genome and the accompanying reads to show that a Prymetime genome assembly is equivalent to the reference using several standard metrics. Finally, we used Prymetime to resequence the nonconventional yeasts *Y. lipolytica* Po1f and *K. phaffii* CBS 7435, producing an improved genome assembly for each strain. Thus, our workflow can achieve accurate, complete, and contiguous whole genome sequences of yeast strains before and after engineering. Therefore, Prymetime enables NGS-based strain quality control through assembly and identification of engineering features.

## Introduction

Whole genome sequencing (WGS) is an attractive method for evaluating genetic engineering because it does not depend on specific sequence features and it captures unintended editing. Yet, engineered organisms are rarely evaluated with WGS, even though the few engineered genomes reported to date show unpredictable features that can only be detected with WGS. These include detection of multiple insertion events^1^, gene loss and chromosomal rearrangement^2^, unexpected mutations affecting phenotype^3^, unpredictable off-target mutations from Cas9 editing^4,5^, insertion of DNA from a plasmid used for cloning^6^, and even insertion of genomic DNA from the cloning host^7^. This does not include a large number of unpublished accounts of WGS revealing unexpected sequences and genome structures in engineered industrial strains. This evidence challenges the assumption that an observed phenotype is the direct result of intended engineering, illuminating a possible explanation for variation between replicates and irreproducible findings - a common problem for biology-related disciplines^8^. Clearly, WGS must be used more broadly to detect and validate genetic engineering.

WGS is particularly needed for engineered yeast strains which can have complex genome features like multiple deletions^9^, multiple plasmids^10^, many insertions^11^, and SCRaMbLEd chromosomes^12,13^. Furthermore, yeast are a crucial testbed for genome-scale design^14,15^, and accurate WGS will be necessary for validating written eukaryotic genomes. Finally, engineered yeast have significant economic value as promising cell factories for the manufacture of medicines^16,17^, fuels^18,19^, materials^20,21^, and chemicals^22,23^. Given the economic importance and increasing use of engineered yeast cell factories, it is crucial that WGS methods are developed that can efficiently validate the presence of intended engineering and confirm the absence of unintended variation. Without practical WGS workflows, the majority of strains are currently validated with inferior methods like PCR and targeted sequencing.

Yet, applying WGS is a challenge because of the diversity of genetic backgrounds, the variety of engineering features, and the current scale of yeast strain engineering. Myriad laboratory strains of the baker’s yeast *Saccharomyces cerevisiae*^9,24,25^ and nonconventional yeasts like *Yarrowia lipolytica*^26–28^ and *Komagataella phaffii* (formerly *Pichia pastoris*)^29,30^ are used to create yeast cell factories, so there are many potential genetic backgrounds. Methods of yeast engineering leave myriad sequence features behind, including standard plasmid sets with standard expression parts^31–34^, high efficiency transformation^35–37^, homologous recombination^10,38–40^, gene knockouts using the Cre recombinase system^41^, and genome editing using RNA-guided endonucleases^7,11,42,44–46^. Furthermore, the scale of yeast engineering is increasing both in the fraction of a genome that may be rewritten^12,47,48^, and in the numbers of engineered strains created through adaptive laboratory evolution^49^-51 and combinatorial pathway engineering^1,52,54-56^. Each of these factors make accurate, complete, and contiguous genomes difficult to attain without significant allocation of resources.

A WGS workflow involves five steps - DNA isolation, sequencing library preparation, sequencing, assembly, and annotation. First, genomic DNA is isolated from all other cellular components using one of a variety of methods, including phenol-choloroform, bead beating, or enzymatic lysis^57^. Second, the sequencing library is prepared by attaching adapters and barcodes. This can be done via ligation, which involves shearing the DNA to create free ends for DNA ligase to attach adapters, or tagmentation, which randomly inserts adapter attachment points without shearing^58^. Third, the library is sequenced with a next-generation sequencing (NGS) platform that either obtains short reads (150-300 base pairs long) with high accuracy58 or long reads (1.5 kilobases to megabases long) with lower accuracy^59^. The average read length and the number of reads (genome coverage) output by the NGS platform is dependent on sequencing technology and the preceding DNA isolation and adapter attachment steps^60^. Fourth, the reads are computationally assembled into a final genome sequence with software that uses either an overlap-layout-consensus (OLC) or De Bruijn graph (DBG) algorithm^61^. OLC algorithms piece together reads based on overlapping sequences to construct progressively larger contiguous sequences (contigs). These algorithms use an All-versus-All consensus step62 that may discard highly identical sequences in order to reach consensus. In contrast, DBG algorithms split reads into shorter k-mers followed by a Eulerian walk approach to construct contigs, thus DBG may be less prone to discarding highly identical sequences^62^. These algorithms assemble a genome sequence *de novo* or, when available, a reference genome may be used to aid assembly^63^. Fifth, an annotation is performed. Eukaryotic annotation involves first predicting genes in the genome sequence, followed by functional annotation^64^. However, engineering features like synthetic biology parts are not annotated. The quality of the assembled, annotated genome sequence is dependent on the read depth, the average read length, and the read accuracy.

Genome assembly quality also depends on genome assembly software. A variety of software based on OLC and DBG algorithms have been made publicly available. These fall into three categories: short read only, hybrid, and long read with error correction. Short read only software uses short read data exclusively, thus it achieves high sequence accuracy but requires high genome coverage to resolve structures like chromosomes as single contiguous sequences (contigs). These include the OLC assembler Edena26 and the DBG assemblers ABySS25 and Velvet^27^. Hybrid assembly software uses both short read and long read data. These assemble short reads first, then stitch contigs together with long reads. Popular hybrid assemblers include the OLC assembler Masurca68 and the DBG assemblers HybridSPAdes69 and Unicycler^28^. Long read with error correction software also uses both short read and long read data. These assemble only long reads with software like the OLC assemblers MiniASM^20^, Canu^22^, and SMARTdenovo23 or the DBG assembler Flye^24^. Due to the error rate of long reads, the resulting contigs are error prone^59^. Therefore, the initial long read contigs only provide a “skeleton” for mapping additional reads^75–84^. The read mapping is performed by error correction software such as Medaka^85^, which uses long read data, and Racon29 or Pilon^31^, which use short read data. Both hybrid and long read with error correction assembly approaches currently hold the most promise to achieve accurate genome sequence and structure at low read depths.

Ideally, one would be able to quickly obtain a genome assembly with WGS that resolves engineering with both accurate sequence and structure (e.g. the correct number of chromosomes and plasmids). Yet, unique challenges arise when applying WGS to engineered yeasts. First, engineered yeasts contain many engineering features that are important to annotate, yet there is currently no way to do this. Second, the high sequence identity in many engineered constructs, such as common plasmid elements or parts derived from the host genome, can cause identical sequences to be omitted^88,89^. In particular, OLC assemblers struggle to reproduce the expected representation and resolution of repeats^20,22,90^. Third, genome assembly software constructs either linear or circular sequences, not both. This is insufficient for yeasts that have a hybrid structure consisting of both linear chromosomes and circular plasmids. Fourth, the scale of yeast strain engineering limits the broad application of WGS due to cost. Currently, iterative design cycles and biological replicates result in many more strains than can be reasonably sequenced. For example, one combinatorial library for itaconic acid production consisted of 1,152 unique strains, including replicates^1^. Another library for penicillin synthesis consisted of 120 unique strains, only a subset of which were validated with targeted sequencing^56^. Therefore, it is necessary for WGS to not only achieve accurate resolution of all engineering signatures, but do so with the minimal amount of resources used per genome.

Here, we present a sample and data processing workflow that is capable of resolving all chromosomes and plasmids within complete, contiguous genomes of engineered yeasts with synthetic biology parts annotated. First, we optimize sequencing library preparation to increase nanopore read length and the number of reads from plasmids. Then, we test different assembly algorithms for their ability to achieve correct, contiguous sequences of the engineering features. We then develop an annotation strategy to label common yeast engineering sequences. We integrate all software steps into a single package, called Prymetime. Then, we validate Prymetime on a panel of 15 engineered yeasts, resolving all engineering sequences in different genetic backgrounds. Further analysis of two strains reveals unintended recombination and insertion events, demonstrating the utility of Prymetime as a quality control tool. We further show that the whole genomes produced by Prymetime using 40X read depth of both nanopore and Illumina reads are equivalent to or better than reference genome assemblies of *S. cerevisiae* CEN.PK113, *Y. lipolytica* Po1f, and *K. phaffii* CBS 7435. Thus, Prymetime allows resolution and annotation of intended engineering signatures, identification of unintended changes, and assembly of quality parent strain genomes.

## Results

### Optimizing Nanopore Sequencing Library Preparation for Engineered Yeasts

From the beginning, we set a standard that our genome assembly workflow must be able to resolve chromosomal integrations and multiple plasmids used in yeast engineering. Therefore, we built a *S. cerevisiae* CEN.PK113 strain containing an integrated carotenoid pathway, the native 2*μ* plasmid, a dCas9 plasmid, and a gRNA plasmid, shown in Figure 1a. We named this strain “FEY_2,” and a picture of several colonies of this strain are shown in 1b. Initially, we prepared sequencing libraries of FEY_2 with a nanopore ligation kit. Sequencing these initial libraries had low average read length that varied from run to run, possibly because of differential DNA shearing during isolation. To limit this, we developed a gentle genomic DNA isolation protocol which increased average nanopore read length and reduced variance (see Supplementary Methods). However, the sequencing results contained very few reads from plasmids, as determined by comparing the average normalized mapped reads of the plasmid antibiotic selection markers to those of the ACT1 genomic locus using Minimap2. We could isolate plasmids from FEY_2 using a yeast miniprep kit, so we reasoned that the sequencing library preparation step was so gentle that it was not linearizing circular plasmids for adapter ligation. Thus, we turned to a tagmentation library preparation method. The improvement in average normalized mapped plasmid reads is shown in Figure 1c for the low copy plasmid and Figure 1d for the high copy plasmid. Interestingly, the 2:1 and 20:1 marker to ACT1 ratios for each plasmid are equivalent to the approximate plasmid copy number in yeast for each origin^25,35^. Furthermore, tagmentation also increased the representation of other circular elements like the native 2*μ* plasmid and mitochondrial DNA. These results indicate that tagmentation is key to achieving long average read lengths while also generating linear molecules from small circular DNA so that they can pass through the nanopore flow cell. Thus, with gentle isolation and tagmentation, nanopore sequencing of FEY_2 resulted in adequate representation of plasmid reads.

**Figure 1.**
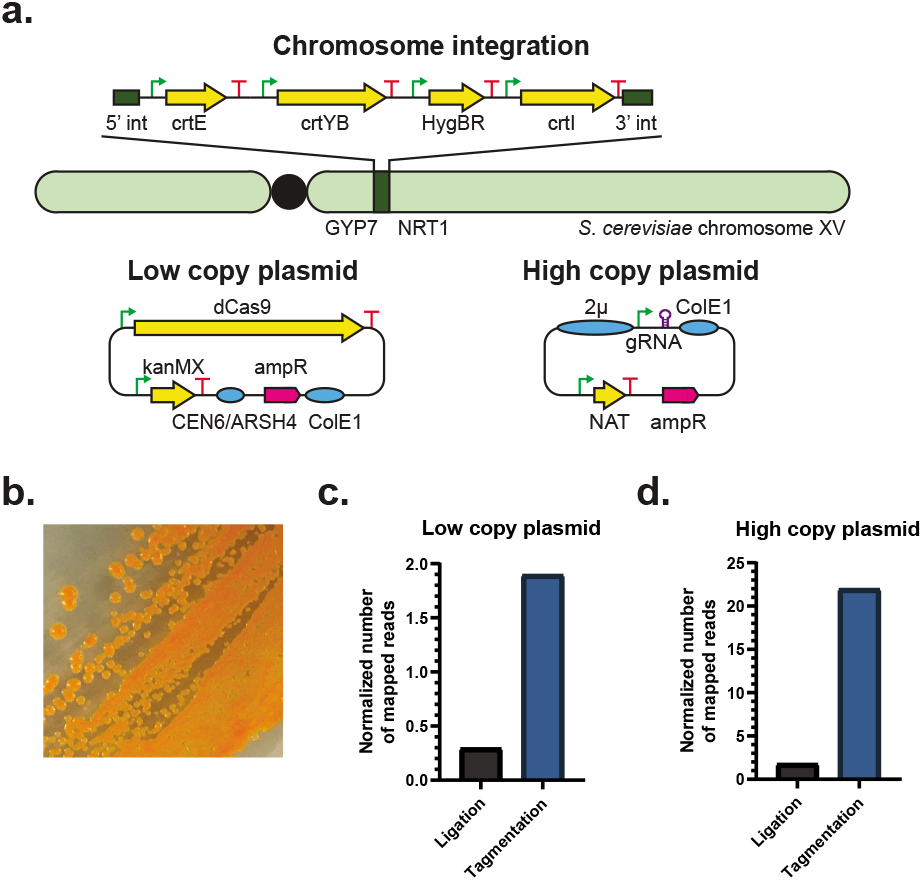
FEY_2 strain design and nanopore library preparation methods affecting FEY_2 read representation. **a.** Illustration of the engineering signatures comprising FEY_2, which included a carotenoid pathway chromosomal integration, a low copy plasmid expressing dCas9, and a high copy plasmid expressing gRNA. **b.** Photograph of FEY_2 streaked onto an agar plate, showing the carotenoid pathway is functional. **c.** Normalized number of mapped reads from libraries prepared by ligation or tagmentation for the low copy plasmid in FEY_2. **d.** Normalized number of mapped reads from libraries prepared by ligation or tagmentation for the high copy plasmid in FEY_2.

### Developing a *de novo* Genome Assembly Workflow for Complete, Contiguous Plasmids and Integrations

Once we achieved appropriate read representation, we investigated which assembly algorithm would correctly assemble the reads into contiguous sequences. This requirement is stringent, particularly for the three plasmids because they each have significant sequence identity between each other and the genome. We evaluated *de novo* assemblers of the following types: short-read only, hybrid, and long read with error correction.

To do this, we used our optimized library preparation to obtain long reads at 60X genome coverage from the ONT MinION and short reads at 125X genome coverage from the Illumina iSeq 100. This common set of reads was then fed to each assembler, and the resulting genome assembly was analyzed using BLASTN for the presence of the integrated pathway, both plasmids, and the native 2*μ* plasmid. A visual representation of the BLASTN results is shown in Figure 2a, with a key in Figure 2b describing the glyphs used to represent the assembly features. The engineering features were rarely complete or assembled into one contiguous sequence. The short-read only *de novo* assemblers ABySS, Edena, and Velvet returned a fragmented, incomplete pathway and plasmids. The hybrid assemblers SPAdes and Masurca produced more complete sequences than the short read only assemblers, but the genome integration was fragmented, and Masurca also omitted portions of the three plasmids. The long read *de novo* assemblers MiniASM, Canu, Flye, and SMARTdenovo each returned a single contiguous sequence for the genome integration, yet, MiniASM, Canu, and SMARTdenovo omitted sections of the three plasmids. Only Flye was able to return the genome integration and each plasmid correctly in contiguous sequences.

**Figure 2.**
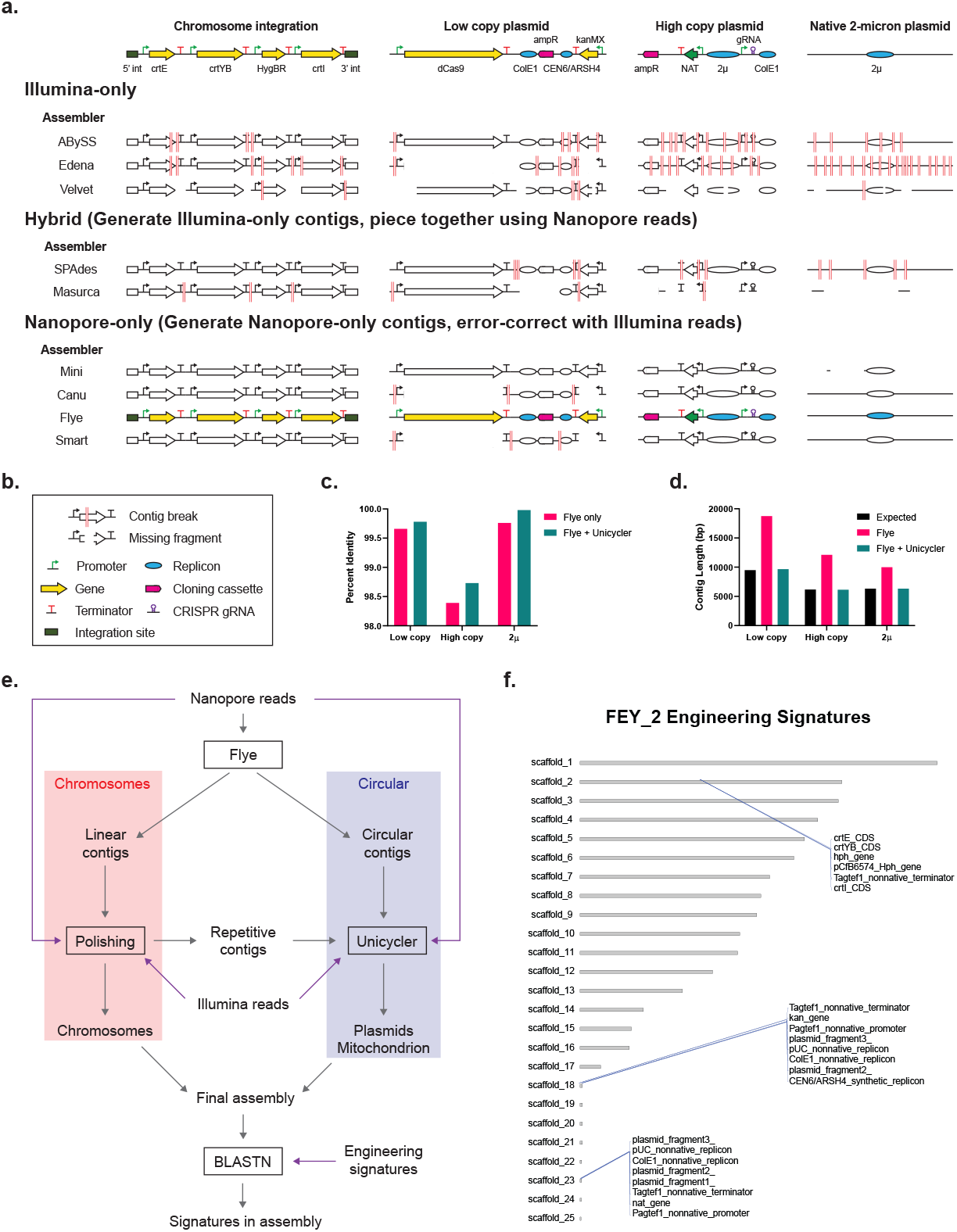
Detection of engineering signatures in S. cerevisiae FEY_2 using various genome assembly strategies. **a.** BLASTN results from querying known engineering signatures against assemblies. **b.** Key describing assembly failure modes and synthetic biology part glyphs. The colored pathways and plasmids represent assemblies where all engineering signatures were found in contiguous sequences. **c.** Plasmid percent identity, found using BLASTN, in the FEY_2 strain both before and after reassembly with Unicycler. **d.** Plasmid contig length in the FEY_2 strain both before after reassembly with Unicycler, compared to the expected plasmid length. **e.** Overview of Prymetime genome assembly pipeline. **f.** Output from Prymetime, showing the FEY_2 genome assembly scaffolds and the engineering signatures found using BLASTN.

Of the assemblies missing large portions of at least one of the three plasmids, almost all were generated with an OLC assembler. To investigate this further, we used BLASTN at each step in the OLC-based Canu pipeline to determine when sequences were omitted. We noted that the complete low-copy plasmid was initially present before the consensus step, and was then lost in the final assembly. It seems that Canu discarded the plasmid at a certain threshold during the consensus step, likely because of high sequence identity with the other plasmids. In contrast, the DBG assemblers Flye, ABySS, and SPAdes did not omit sections of plasmids. While the plasmids were fragmented among different contigs with ABySS and SPAdes, Flye assembled all plasmids into single contiguous sequences. This indicates that, along with sequencing library preparation, genome assembly quality is also dependent on the *de novo* assembler.

While the plasmid contigs from Flye were complete, they were longer than expected. Further inspection revealed that the contigs consisted of several repeats of the expected plasmid sequence. This is a common problem for long-read assemblers, as they use linear logic to merge contigs^22,91^. To obtain structurally representative plasmid contigs, we wrote a script to identify putative circular sequences and send them to be re-assembled with Unicycler, software that was built to assemble circular contigs from bacterial sequencing data^28^. Reassembly of plasmids with Unicycler improved the accuracy as measured by BLASTN (Figure 2c), and length (Figure 2d) of the contigs for the three plasmids in FEY_2.

To simplify the search for engineering features with BLASTN, we added a step that annotates features in the genome from synthetic biology part sequences. We developed a list of many yeast engineering features, including promoters, terminators, selection markers, fluorescent reporters, common coding sequences, origins, and conserved plasmid fragments. This is the default part annotation list for Prymetime, but the user may easily use a different list. A sequence in the genome assembly with high identity to a part on the list are annotated and then plotted using the genome plotter karyoploteR^36^, shown in Figure 2f for the FEY_2 genome assembly. This feature allows the user to quickly visualize all engineering signatures in the entire genome, particularly highlighting plasmids and integrations. Additionally, this feature permits identification of known engineering sequences in an unknown sample.

We coded the final workflow into a single dockerized software package called Prymetime: “Pipeline for Recombinant Yeast genoMEs That Identifies Markers of Engineering.” To our knowledge, this is the first workflow able to annotate engineering features in yeast genomes and assemble both linear and circular contigs. An overview of the workflow is shown in Figure 2e, with a more detailed diagram in Figure S1. In our automated workflow, Flye first assembles nanopore reads and classifies contigs as linear or circular. Then, linear contigs are error-corrected using the polishing software Medaka, Racon, and Pilon, while the circular contigs are re-assembled with long read and short read data using Unicycler. Finally, in an optional step, engineering features are annotated and visualized with karyoploteR. As Figure 2f shows, this workflow assembled chromosomes, plasmids, and successfully annotated engineering features in the FEY_2 genome.

### Resolving Engineering Signatures in a Collection of Engineered Yeasts

We next validated Prymetime on a collection of engineered laboratory and nonconventional yeast. We constructed 15 strains from *S. cerevisiae* S288C, CEN.PK113-7D, W303-*α*, BY4741, BY4742, and *K. phaffii* ATCC 76273 (CBS 7435)93'94 and *Y. lipolytica* ATCC MYA-2613 (Po1f)^95^. A description of each strain is shown in Figure 3, with more detailed descriptions of each strain in Table S1. Engineering signatures were inserted into the genome or maintained on episomal plasmid*S. S. cerevisiae* integrations were targeted to the HO locus32 or between NRT1 and GYP7 in chromosome XV^1,45^. *S. cerevisiae* plasmids consisted of custom TypeIIS-compatible yeast shuttle vectors with either *S. cerevisiae* replicon (2*μ* or CEN6/ARSH4). Engineering was broadly categorized into biosynthetic pathways, gene editing components, deletions, and synthetic biology elements. Biosynthetic pathways included propane^96^, *β*-carotene^6^, prespatane^10^, carnosic acid^13^, and limonene^100,101^. Genome editing associated tools included SpCas9^8^, dCas^97^, LbCpf1^45^, FnCpf1^44^, and Cre recombinase^41^. Deletions included the synthetic auxotrophies already present in *S. cerevisiae* W303-*α*, BY4741, BY4742, and *Y. lipolytica* Po1f. Synthetic biology elements included fluorescent proteins^9^,16 and the 2A sequence^17^. The engineered *Y. lipolytica* strain “FEY_74” contained a CRISPR-Cas9 expression plasmid that contained a codon-optimized version of the Cas9 protein, along with a gRNA expression cassette^46^. The engineered *K. phaffii* strain “FEY_75” contained a chromosomally-integrated red fluorescent protein (RFP) cassette. This strain was transformed using a two-step recombinase based system with integrative plasmids^34^.

**Figure 3.**
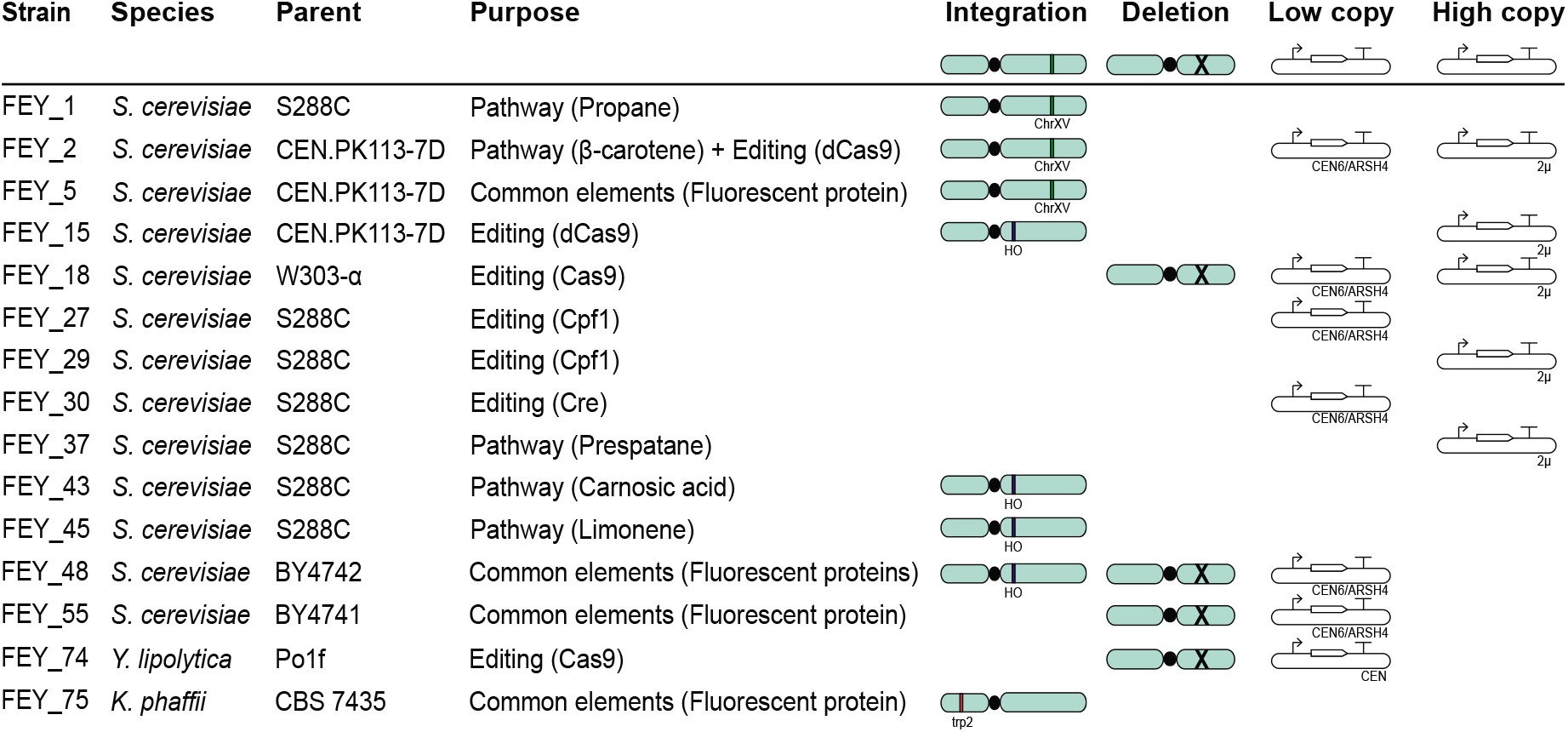
Panel of diverse engineered yeast strains constructed. Presence of an integration, deletion, low copy plasmid, or high copy plasmid icon indicates a strain has the respective engineering signature, while the absence of an icon indicates it does not have the engineering signature.

We sequenced this collection with the ONT MinION and the Illumina iSeq 100. From the sequencing data, Prymetime produced genome assemblies that captured each engineering signature in each *S. cerevisiae* genetic background as measured by BLASTN of the reference sequence against the assembly. Shown in Figure 4a, the genome assemblies resolved seven different genome integrations in two genome loci and eleven different plasmids. Metrics for the BLASTN results for all engineered *S. cerevisiae* strains are in Table S2, while the annotated karyoploteR visualizations can be found in Figure S2. Figure S3 shows that including Unicycler for plasmid assembly improves length and accuracy in every strain, not just FEY_2. Furthermore, neither the type of gene (metabolic, selective, editing, or reporter), nor repetitive parts (Ptef1, Pgal10), nor plasmid copy number affected the accuracy or structural completeness of the assembly.

**Figure 4.**
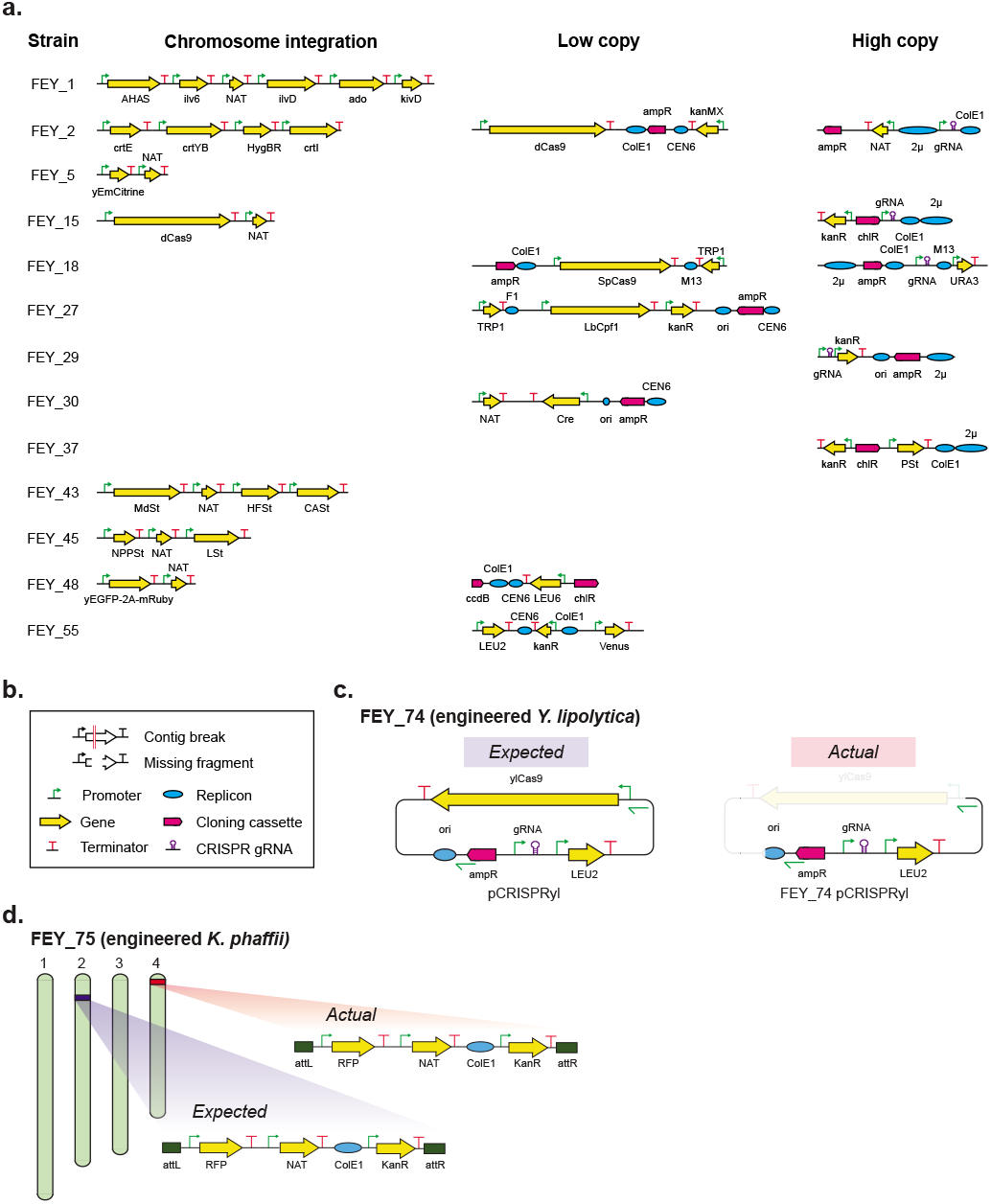
Resolving signatures of engineering from the panel of engineered yeast strains. **a.** Visual representation of the BLASTN results from querying known engineering signatures against Prymetime-assembled genome assemblies of the 15 engineered *S. cerevisiae* strains. **b.** Key describing assembly failure modes and synthetic biology part glyphs. The colored pathways and plasmids represent assemblies where all engineering signatures were found in contiguous sequences. **c.** The expected CRISPR-Cas9 expression vector for FEY_74, an engineered *Y. lipolytica* strain, and the actual plasmid from the Prymetime genome assembly (see Figure S3). **d.** Illustration showing the expected location of the RFP integration cassette into chromosome II of FEY_75, an engineered *K. phaffii* strain, and the actual location of the cassette into chromosome IV.

The genome assemblies from the two engineered nonconventional yeasts - *Y. lipolytica* strain FEY_74 and *K. phaffii* strain FEY_75 - revealed unintentional engineering. FEY_74 was intended to contain the pCRISPR-yl46 plasmid, shown in Figure 4c. However, the plasmid contig from the genome assembly was missing the entire Cas9 transcription unit and a portion of the *E. coli* origin of replication. Inspection of the raw reads failed to identify a single read with the missing ylCas9 sequence. We performed a genomic DNA isolation and a yeast plasmid miniprep on FEY_74 and transformed the resulting DNA back into *E. coli*, yet did not observe any colonies. This indicates that the disrupted origin of replication in the assembly reflects an actual unintended loss rather than an assembly error. This was further confirmed by PCR of DNA isolated from FEY_74 with primers spanning the missing region of the plasmid. The length of the PCR product indicated that the Cas9 transcription unit was indeed missing (Figure S4). Similarly, FEY_75 was designed to have an RFP transcription unit integrated into chromosome II (Figure 4d). However, PCR of the integration site in chromosome II failed to confirm integration, even though the strain was nourseothricin resistant and RFP positive. Prymetime was able to annotate the entire pathway in the FEY_75 genome, but the pathway was integrated into chromosome IV. These results indicate that Prymetime can be used to find and validate engineering sequences, which is useful for both strain quality control and identification of engineering in unknown samples.

### Whole Genome Assembly Quality

We then analyzed the whole genome quality of the Prymetime assemblies. Each engineered *S. cerevisiae, Y. lipolyitica*, and *K. phaffii* genome had high contiguity, sequence accuracy, and genome completeness. However, to clearly show that Prymetime can assemble quality whole genomes we benchmarked it against the best publicly available *S. cerevisiae* CEN.PK113-7D genome assembly for which the raw reads were also available^105,106^. We used these same raw reads to generate a genome assembly with Prymetime. As shown in Figure 5a, the reference assembly used raw nanopore, Illumina, and PacBio reads^106^, while Prymetime assembled random subsets of the raw nanopore and Illumina reads at different read depths. These subsets were passed through each step in Prymetime (Figure 5b). We compared the assembly quality at each step to the reference genome.

First, we evaluated the structure of the initial contigs from Flye using metrics from QUAST^32^, specifically, N50, the number of contigs, and the length of the largest contig (Figure 5c). The standard deviation was calculated from the variation in the metrics resulting from three different random read subsets. The Flye step achieves N50 equivalent to the *S. cerevisiae* CEN.PK113-7D reference at a long read genome coverage of 40X and above. Similarly, 40X genome coverage and above produced the expected 18 contigs - sixteen chromosomes, the native 2*μ* plasmid, and mitrochondrial DNA. Further, the longest contig from 40X genome coverage and above is equivalent to the reference genome. To be thorough, we also tested nanopore assemblers other than Flye to see if these produced improvements, depicted in Table S3. Flye remained the best assembler. These results indicate that the Flye assembly step can generate reference quality genome structure with a minimum long read sequencing depth of 40X.

Next, we evaluated the sequence accuracy of Prymetime. Average identity to the reference genome was calculated with MUMmer33 at different points in the Prymetime polishing workflow, shown in Figure 5d. The unpolished long read assembly from Flye only matches 98.1% of the reference genome, while successive Medaka (long read polishing), and Racon and Pilon (short read polishing) steps improve the assembly, eventually matching the reference. Then, we assessed the read depth of short reads needed to optimize Racon and Pilon polishing. To do this, we calculated average identity to reference and the number of single nucleotide polymorphisms (SNPs) for different short read genome coverages, again using MUMmer. The results indicate that short reads at 40X genome coverage and above are sufficient to match the reference genome. Additionally, the accuracy of polishing assemblies from each nanopore assembler other than Flye is presented in Table S4 (adding polishing steps) and Table S5 (changing short read depth). These results indicate that successive polishing with Illumina short reads at a minimum read depth of 40X is critical for sequence accuracy.

Finally, the polished genome assemblies were evaluated for completeness. Two completeness metrics were used - the percentage of *S. cerevisiae* S288C open reading frames (ORFs) contained in an assembly, calculated by BLASTN, and the Benchmarking Universal Single-Copy Orthologs (BUSCO) score, calculated using the Saccharomycetales dataset^34^. The results are shown in Figure 5e. In terms of percentage of *S. cerevisiae* S288C ORFs, Prymetime genome assemblies using a long read depth of 40X most closely matched the *S. cerevisiae* CEN.PK113-7D reference. In terms of the BUSCO score, all assemblies with long read depths above 20X were equivalent to the reference. Additionally, these metrics were calculated for polished assemblies using other nanopore assemblers, shown in Table S5 (BUSCO score) and Table S6 (percent of S288C ORFs). Taken together, these results indicate that the genome assemblies generated by Prymetime are structurally correct, accurate, and complete using only 40X read depth for both long and short reads.

**Figure 5.**
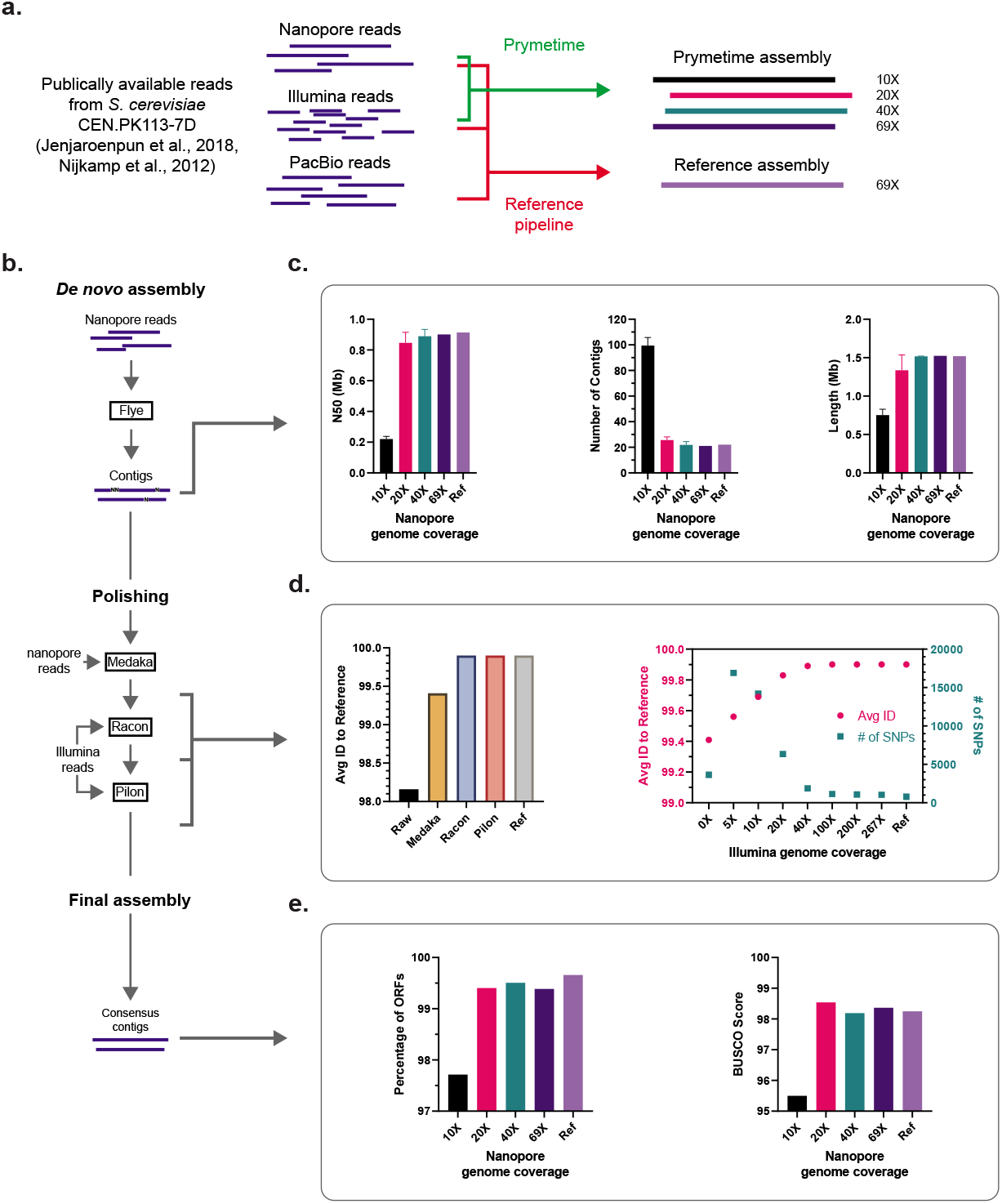
Genome assembly quality at varying genome coverage depths using publicly available reads from a S. cerevisiae CEN.PK113-7D reference genome. **a.** Prymetime and reference assembly workflows and read depths. **b.** Assembly and polishing workflow. Assemblies were evaluated at different points in this workflow, indicated by the arrows. **c.** Genome structure comparison, using QUAST. **d.** Genome accuracy with different polishing steps and read depths. **e.** Genome completeness quantified by the percentage of open reading frames (ORFs) from *S. cerevisiae* S288C in each genome assembly and by the BUSCO score.

### Re-Sequencing Nonconventional Yeast

To demonstrate that the entire laboratory and computational workflow can obtain high quality reference genomes, we re-sequenced the two nonconventional yeasts in this study with the ONT MinION and Illumina iSeq 100. The resulting *de novo* genome assemblies output from Prymetime were compared to the publicly available reference assembly for *Komatagaella phaffii* CBS 7435110 and *Yarrowia lipolytica* Po1f^28^. Comparing the whole genomes with Mauve111 qualitatively confirmed the completeness of the Prymetime assemblies (Figure 6a depicts *K. phaffii* and Figure 6b depicts *Y. lipolytica*). Interestingly, Prymetime resolved a large region in the third contig of *Y. lipolytica* Po1f that is not in the reference (purple shading in Figure 6b). Quantitatively, the Prymetime genome assemblies had high genome contiguity as measured by the number of contigs (Figure 6c), had no assembly gaps (Figure 6d), and improved genome completeness measured by BUSCO score (Figure 6e). The higher BUSCO scores are because there are 6 more essential genes in the *K. phaffii* Prymetime assembly and 13 more essential genes in the Y *lipolytica* Prymetime assembly. This is, to our knowledge, the first assembly of these essential genes in these nonconventional yeasts. Overall, these results indicate that Prymetime can be used to generate high quality *de novo* reference genomes of nonconventional yeasts.

**Figure 6.**
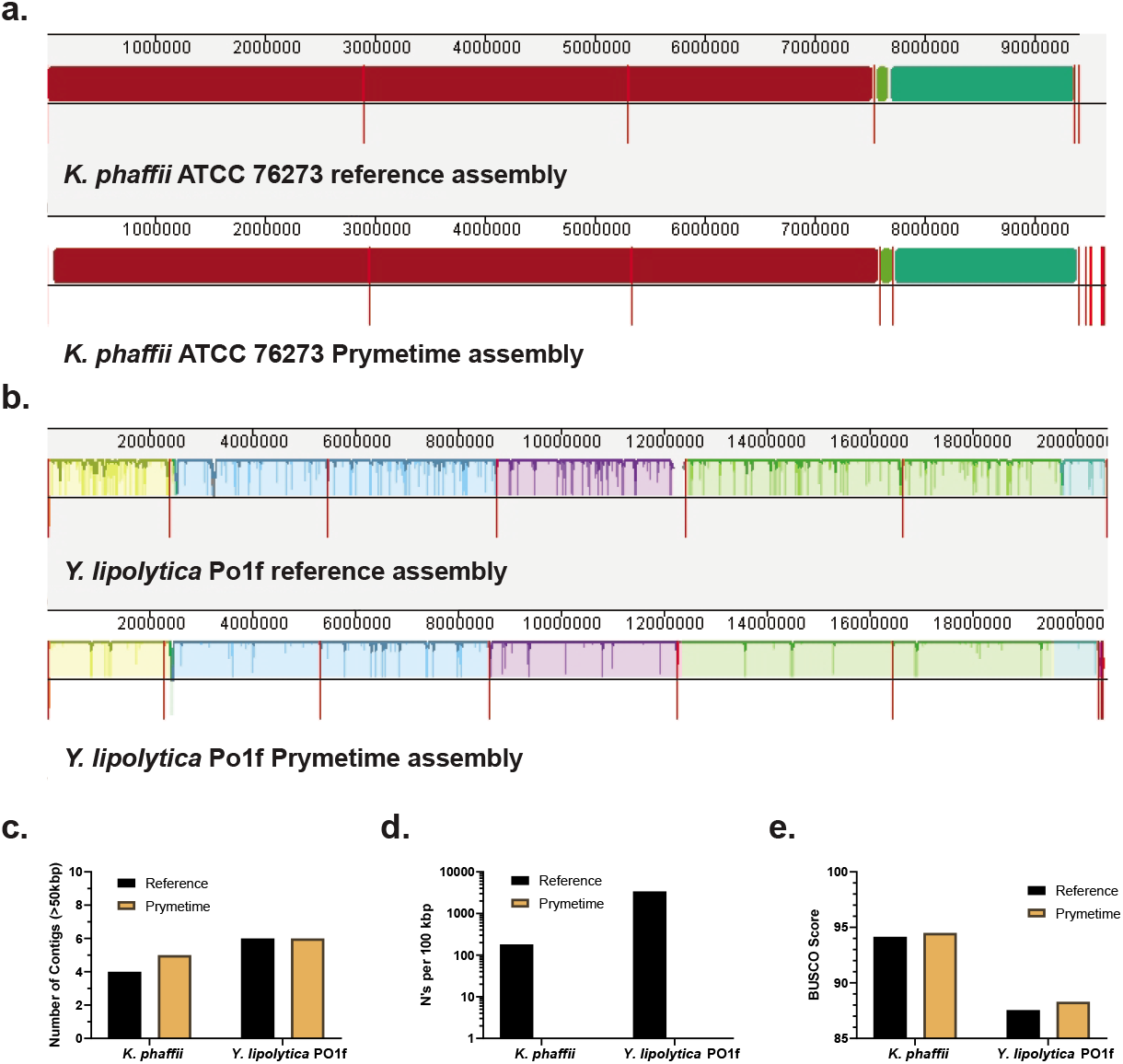
Re-sequencing of two nonconventional yeast strains. **a.** Mauve visualization of a *K. phaffii* whole genome comparison between the publicly available reference assembly and the Prymetime assembly. **b.** Mauve visualization of a Y *lipolytica* whole genome comparison between the publicly available reference assembly and the Prymetime assembly. The red lines on each alignment indicate a new contig. **c.** Number of contigs in the reference genome assemblies versus the Prymetime genome assemblies. **d.** Gaps in the reference genome assemblies versus the Prymetime genome assemblies. The number of gaps were represented by the number of N’s per 100 kbp. **e.** Genome BUSCO score.

## Discussion

This work describes development of a novel workflow for WGS of engineered yeasts. The validated workflow consists of gentle gDNA isolation, tagmentation, long and short read NGS, accurate *de novo* assembly of both linear and circular elements, and synthetic biology part annotation. Using this workflow, diverse engineering signatures can be resolved in complete, contiguous sequences even with multiple similar plasmids in one strain. The resulting whole genome quality is comparable to high-quality reference assemblies, therefore, it is possible to generate accurate genome assemblies both before and after engineering. This permits verification of genetic engineering in yeasts with WGS.

Interestingly, only Flye was suitable for the first assembly step, fully satisfying the strict requirements of engineering signature accuracy, completeness, and contiguity. The other *de novo* genome assemblers we tested omitted engineering signatures at the read depths tested - even with appropriate representation of plasmid reads. This is because the strain used to benchmark the assemblers had multiple plasmids with high sequence identity between them. We observed that signature omission most commonly occurred with assemblers built around OLC algorithms. Assemblers built around DBG algorithms were consistently better at resolving all signatures. Yet, Flye remained the only assembler able to resolve all of the signatures in contiguous sequences. These observations highlight the difficulty of applying otherwise effective genome assembly software to engineered yeasts, which have complex and repetitive sequence elements, and the need to continually improve assembly algorithms to better handle complex features. Based on our results, we recommend benchmarking future assemblers on the performance of Flye.

Whole genome sequencing is rarely used in strain engineering cycles due to the barriers of NGS cost, time, and required bioinformatics expertise. The WGS workflow we developed with the inexpensive ONT MinION and Illumina iSeq 100 platforms and the integrated, dockerized Prymetime software package overcomes these barriers. With Prymetime, we were able to achieve high-quality genomes at relatively low read depth, finding that 40X for both long and short reads was sufficient for accuracy, completeness, and contiguity of whole yeast genomes and the engineering features within them. With 40X read depth, up to 30 *S. cerevisiae* genomes can be sequenced on one MinION flow cell and up to 4 genomes can be sequenced on one Illumina iSeq flow cell. This is because 0.5 Gb is needed for 40X read depth of the 12.1 Mb *S. cerevisiae* genome and our typical yield is approximately 15 Gb from the MinION and 2.4 Gb from the iSeq 100. Not accounting for labor, this level of multiplexing would cost around $200 per genome. The entire workflow is fast - it takes under a week to start from a single colony and acquire a genome assembly, requiring only 15 hours of hands-on time. Finally, our workflow requires only a few coding steps - future users can simply load NGS reads and run the Prymetime script (see the GitHub repository at https://github.com/emyounglab/prymetime for more details).

The integrated workflow described here permits rapid, on-site acquisition of reference quality yeast genome sequences and annotation of genetic parts. Thus, it detects and validates genetic engineering in yeasts. The utility of this workflow for verification of engineering and resequencing of nonconventional yeasts was demonstrated, indicating that it may also be applied to sequence novel yeast isolates and identify engineering in unknown samples. We envision that this novel approach is broadly applicable to any effort that involves engineered yeasts.

## Data Availability

The reference genomes for *S. cerevisiae, Y. lipolytica*, and *K. phaffii* can be accessed on NCBI. All engineered genome sequences are available on NCBI. The raw reads have been deposited on NCBI.

## Code Availability

Prymetime can be accessed as a Docker image on GitHub at https://github.com/emyounglab/prymetime.

## Acknowledgements

The authors thank James Kingsley at WPI for his help implementing Prymetime on WPI’s server. This research is based upon work supported in part by the Office of the Director of National Intelligence (ODNI), Intelligence Advanced Research Projects Activity (IARPA) under Finding Engineering Linked Indicators (FELIX) program contract #N66001-18-C-4507. The views and conclusions contained herein are those of the authors and should not be interpreted as necessarily representing the official policies, either expressed or implied, of ODNI, IARPA, or the U.S. Government. The U.S. Government is authorized to reproduce and distribute reprints for governmental purposes notwithstanding any copyright annotation therein. This work is also supported by Worcester Polytechnic Institute startup funds.

## Author Contributions Statement

JHC, AA, NR, and EMY conceived of the study. JHC conducted all sequencing runs and bioinformatics analysis. JHC, KWK, TRJ, SB, CBM, MC, and ZJN built the collection of engineered yeasts for sequencing. JHC, TM, and EMY wrote PRYMETIME scripts and created the Docker image.

