## Supplemental_Information for "Verification of genetic engineering in yeasts with nanopore whole genome sequencing"

#### Table of Contents

Supplementary Methods

Table S1: Detailed descriptions of engineered strains.

Table S2: BLASTN results for all engineered signatures in their respective genome assemblies.

Table S3: Evaluation of *de novo* Nanopore assemblers at various genome coverage depths with structure-related metrics.

Table S4: Improvement to assembly accuracy through polishing at various genome coverage depths of Illumina reads.

Table S5: Improvement to assembly accuracy using different polishing tools.

Table S6: BUSCO genome completeness assessment for each Nanopore *de novo* assembler at each genome coverage depth and polishing software.

Table S7: Percentage of *S. cerevisiae* S288C CDS found in each Nanopore *de novo* assembler at each genome coverage depth and polishing software.

Figure S1: Comparison of engineered strain assembly plasmids with or without the added Unicycler step.

Figure S2: Genome plots of each engineered strain and their associated non-native engineering signatures.

Figure S3: Detailed Prymetime genome assembly pipeline.

Figure S4: DNA gel of a PCR across the dCas9 gene of the pCRISPRyl plasmid from the FEY_74 strain

Figure S5: Whole genome comparison of engineered *S. cerevisiae* strains against their respective parent strains using Mauve.

Figure S6: Whole genome comparison of engineered nonconventional strains against their respective parent strains using Mauve.

### Supplementary Methods

#### Microbial Culture

Yeast strains were grown in yeast extract-peptone-dextrose (YPD) or synthetic complete (SC)+glucose media. YPD consisted of 30 g/L YEP (10 g/L yeast extract + 20 g/L peptone, Sunrise Science, 1877-1KG) and 20 g/L glucose (Alfa Aesar, A16828). SC+glucose media consisted of 6.71 g/L of YNB+Nitrogen (1.71 g/L yeast nitrogen base + 5 g/L ammonium sulfate, Sunrise Science, 1501-250), 20 g/L glucose, and a formulation of complete synthetic media (CSM). CSM formulations were 1) CSM-Leu: 0.65 g/L CSM-His-Leu-Ura (Sunrise Science, 1015-010) + 0.02 g/L Histidine (Sunrise Science, 1978-010) + 0.02 g/L Uracil (Sunrise Science, 1906-010) and 2) CSM-Trp-Ura: 0.62 g/L CSM-Leu-Trp-Ura (Sunrise Science, 1017-010) + 0.1 g/L Leucine (Sunrise Science, 1980-010). If appropriate, antibiotic selection was performed with nourseothricin at 0.1 g/L for *S. cerevisiae* and *K. phaffii* (Jena Bioscience, AB-101-10ML), geneticin at 0.2 g/L for *S. cerevisiae* and 0.3 g/L for *K. phaffii* (Life Technologies Gibco, 10131-035), and/or hygromycin B at 300 mg/L for *S. cerevisiae* (Thermo Fisher, 10687010).

Chemically competent *E. coli* DH5$\alpha$ (NEB, C2987H) was used as a cloning strain and grown in 25 g/L LB Miller broth (10 g/L tryptone + 5 g/L yeast extract + 10 g/L sodium chloride, Fisher Scientific, BP1426-2). Antibiotic selection was performed with 100 mg/L ampicillin (Alfa Aesar, J63807), 25 mg/L chloramphenicol (Alfa Aesar, B20841), or 50 mg/L kanamycin (Alfa Aesar, J61272). Solid media was supplemented with 20 g/L agar (Sunrise Science, 1910-1KG).

#### Strain Design and Construction

Gene expression parts for *S. cerevisiae* were sourced from a previous study[1]. The parts were maintained and combined using the hierarchical Type-IIS assembly system from this study. The MoClo Pichia Toolkit (Addgene, Kit #1000000108) was used to construct the expression vector for *K. phaffii*[2]. All enzyme genes were designed and synthesized by Integrated DNA Technologies (IDT). Design included codon optimization using IDT's proprietary algorithm and elimination of BsaI and BbsI restriction sites. The sequences for acetolactate synthase(ahas1), ketol-acid reductoisomerase (ilv6), and dihydroxy-acid dehydratase (ilvD1) were derived from *Penicillium chrysogenum*[3]. The sequence for aldehyde decarbonylase (ado) was derived from *Prochlorococcus marinus*[4]. The sequence for alpha-ketoisovalerate decarboxylase (kivD) was derived from *Lactococcus lactis*[5]. The sequences for geranylgeranyl diphosphate synthase (crtE), bifunctional lycopene cyclase/phytoene synthase (crtYB), and phytoene desaturase (crtI) were sourced from a previous study[6]. The deactivated Cas9 (dCas9Mx1) sequence was sourced from a previous study[7]. The gRNA cassette was sourced from a previous study[8]. The sequence for the fluorescent protein encoding gene yEmCitrine was sourced from a previous study[9]. The sequence for prespatane sesquiterpene synthase (pst) was derived from *Laurencia pacifica*[10]. The sequence for bifunctional diterpene synthase (mdst) was derived from *Selaginella moellendorffii*[11]. The sequence for bifunctional ferruginol, 11-hydroxyferruginol synthase (hfst) was derived from *Salvia pomifera*[12]. The sequence for 11-hydroxyferruginol C20-oxidase (cast) was derived from *Salvia rosmarinus*[13]. The sequence for dimethylallylcistransferase (nppst) was derived from *Solanum lycopersicum*[14]. The sequence for limonene synthase was derived from *Citrus limon*[15]. The yEGFP-2A-mRuby sequence was designed by combining the yEGFP and mRuby sequences from Sheff et al.[9] and Lee et al.[16], respectively, with a self-cleaving 2A sequence[17]. The sequence for b-isopropylmalate dehydrogenase (Klleu2) was sourced from a previous study[18]. The sequence for the Venus gene was sourced from a previous study[1].

#### High-molecular weight genomic DNA isolation

Genomic DNA was isolated using Promega's Genomic DNA Isolation Kit (Promega, A1120). A modified version of Promega's protocol for yeast gDNA isolation was used to limit shearing of DNA, with added insight from Josh Quick's Ultra-long read sequencing protocol[19]. No vortexing and limited pipetting/mixing steps were used to maximize Nanopore read lengths. 5 mL of cells were grown overnight (or until saturation) at 30°C. The cells were pelleted at 500 x g for 5 minutes, and resuspended in 1.5 mL of 50 mM EDTA and 37.5 $\mu$L of 5 U/$\mu$L zymolyase (Zymo, E1004). The samples were incubated at 37°C for 1 hour to allow the Zymolyase to digest the cell wall. The cells were pelleted at 500 x g for 5 min, re-suspended in 1.5 mL of the Nuclei Lysis Solution (mix by inversion, flicking), and incubated at room temperature for 30 minutes. 7.5 $\mu$L of RNAse A Solution was then added and incubated for 15 minutes at 37°C. Once cooled to room temperature, 500 $\mu$L of the Protein Precipitation Solution was added (invert to mix). The samples were put on ice for 5 minutes, and subsequently centrifuged for 10 minutes at 3000 x g. 700 $\mu$L of the supernatant was added to a fresh microcentrifuge tube with 700 $\mu$L of isopropanol. The microcentrifuge tubes were gently mixed by inversion and centrifuged at 4000 x g for 1 minute. The DNA pellet was washed with 70% ethanol and centrifuged at 4000 x g for 1 minute. The ethanol was carefully pipetted off the DNA pellet, and the tube cap was left open at room temperature for 20 minutes to allow residual ethanol to evaporate. 50 $\mu$L of 10 mM Tris-HCl and 0.02% Triton X-100 was added to resuspend the DNA pellet and incubated overnight at 4°C. DNA quality was evaluated using a Nanodrop, and the concentration was calculated using a Qubit.

##### Nanopore DNA Library Preparation and MinION loading

The Rapid Barcoding Kit was used to tagment the DNA libraries for sequencing (ONT, SQK-RBK004). Up to four genomes were multiplexed on each MinION flow cell. Library preparation closely followed the protocol provided by ONT. Briefly, 400 ng of template DNA for each isolate was diluted to 7.5 $\mu$L, mixed with 2.5 $\mu$L of the Fragmentation Mix, and then incubated at 30°C for 1 minute and 80°C for 1 minute on a thermal cycler. The barcoded samples were then pooled together and concentrated using AMPure XP beads in 10 $\mu$L of 10 mM Tris-HCl, 50 mM NaCl. The pooled sample was next mixed with 1 $\mu$L of RAP for 5 minutes at room temperature, and stored on ice until ready to load. R9.4 MinION Flow Cells (ONT, FLO-MIN106) were used for all sequencing runs. The flow cells were first primed per ONT's instructions. The 11 $\mu$L of prepped DNA was mixed with 4.5 $\mu$L of nuclease-free water, 34 uL of SQB and 25.5 $\mu$L of LLB, and loaded onto the MinION flow cell. Sequencing runs were executed using ONT's MinKNOW software with the default settings.

##### Read processing

Nanopore fast5 files were basecalled using Guppy v2.3.5 (Oxford Nanopore base caller). The subsequent fastq files were demultiplexed using the EPI2ME interface (Metrichor, Oxford, UK). Illumina reads were demultiplexed using the native software on the iSeq machine. Random subsets of Illumina and Nanopore reads at a specific genome coverage were generated using a custom python script (https://github.com/aseetharam/common_scripts/blob/master/sample_fastq.py).

##### Illumina DNA Library Preparation and iSeq 100 loading

The Nextera DNA Flex Library Prep Kit (Illumina, 20018704) along with the Nextera DNA CD Indexes (Illumina, 20018707) were used to tagment the DNA libraries for sequencing. Library preparation closely followed the instructions provided by Illumina, and up to four genomes were multiplexed on one Illumina sequencing cartridge. Briefly, tagmentation was first performed with 500 ng of genomic DNA in 30 $\mu$L of nuclease-free water. The reaction was stopped by adding 10 $\mu$L of both the BLT and TB1 reagents and incubating at 55°C for 15 minutes on a thermal cycler. Index adapters for each sample along with EPM were then added to barcode and amplify the genomic DNA. The DNA libraries were amplified using the following PCR program:

1. 68C for 3 min
2. 98C for 3 min
3. 5 PCR cycles:
   (a) 98C for 45 sec
   (b) 62C for 30 sec
   (c) 68C for 2 min
4. 68C for 1 min
5. 10C hold

The DNA libraries were cleaned using subsequent steps with the SPM reagent and 80% ethanol, and concentrated in 32 $\mu$L of the RSB reagent. Assuming four genomes were multiplexed on one flow cell, 25 pM of each DNA library were pooled together in 100 $\mu$L of the RSB reagent and stored on ice until ready to load. The pooled libraries were loaded onto the sequencing cartridges according to Illumina's instructions. The Local Run Manager on the iSeq 100 machine was used to initiate sequencing runs. A GENERATEFASTQ run was started, and run with the parameters Read Type: Paired End, Read Lengths: 151, and Index Reads: 2.

##### Nanopore *de novo* genome assembly

For the MiniASM[20] assembly, reads were first mapped using minimap2[21] with the parameters "-x ava-ont -t8". MiniASM (v0.3) was then subsequently run with the default parameters. Canu[22] (v1.8) was run with the parameters "minReadLength=2500 mhapSensitivity=high corMhapSenstivity=high corOutCoverage=500". SMARTdenovo[23] (v1.0) was run using the parameters "-c 1 -k 14 -J 2500 -e zmo". Flye[24] (v2.4) was run with the parameters "–meta –plasmids". ABySS[25] (v 2.1.5) was run with the abyss-pe option and the parameter "k=96". Edena[26] (v3.131028) was run with the default parameters. Velvet[27] (v1.2.10) was run with a hashlength of 21 bp. MaSuRCA (v3.3.4) was run with the parameter "JF_SIZE = 242000000 FLYE_ASSEMBLY=1". SPAdes (v3.13.1) was run with the parameters "–sc –nanopore –pe< # >-1 –pe< # >-2". Unicycler[28] was run with the default parameters for using both Illumina and Nanopore reads.

##### Nanopore and Illumina read polishing

The *de novo* Nanopore genome assemblies were first polished with Nanopore reads using Medaka (v0.4) with the default parameters. The assembly was then polished with Illumina reads, first with Racon followed by Pilon. For Racon, the Illumina reads were first mapped to an assembly using minimap2 with the parameters “-ax sr”. Racon[29] was then run using the default parameters. For Pilon, assemblies were first indexed using bwa[30]. Illumina reads were then mapped to the assembly using bwa with the parameters “mem –t 14”. Pilon[31] was then run using the parameter “-Xmx160G”.

##### Genome assessment tools

QUAST[32] (v5.0.0) was run with the default parameters, yielding the metrics number of contigs, maximum contig length, and N50. For accuracy related metrics, the nucmer command was run as part of the MUMmer package[33]. The command "dnadiff -d" was used on the resulting delta file to find the average identity to the reference and the number of SNPs. Genome assemblies were evaluated for genome completeness using BUSCO[34] with the saccharomycetales_odb9 datasets, as well as a BLASTN[35] search of ORFs from *S. cerevisiae* S288C. Engineered signatures were searched for in genome assemblies using BLASTN with the expect threshold set at 0.0001.

##### Detection of non-native engineering signatures

Non-native engineering signatures were detected in the genome assemblies using BLASTN. A curated list of known engineering signatures in fasta format was input to Prymetime using the -eng_sig option. This list contains all non-native engineering signatures used to engineer the yeast strains, and can be downloaded from the Prymetime github page. The genome plotter karyoploteR[36] was used to show the engineering signature hits from the BLASTN search in the context of the entire genome assembly.

#### Supplementary Tables

**Table S1.** Detailed descriptions of engineered strains

| Strain | Parent | Knockout(s) | Chromosome Insert | Plasmid 1 | Plasmid 2 | Plasmid Source |
|---|---|---|---|---|---|---|
| FEY_1 | *S.c.* S288C | - | hap1:HAP1 ChrXV::(Ptef1-ahas1-Ttip1//Psmtef1-ilv6-Tprm9//Phta1-ilvD1-Tyhi9//Pagtef1-nat-Tagtef1//Psptdh3-ado-Trpl41b//Ptdh3-kivD-Trpl15a) | - | - | - |
| FEY_2 | *S.c.* CENPK | - | ChrXV::(Psbtdh3-crtE-Trpl41b//Psptdh3-crtYB-Tyol036w//Phta2-hyg-Tagtef1//Psmtef1-crtI-Trpl15a) | pAG700::(Ptdh3-dCas9-Tadh1//Pagtef1-nat-Tagtef1//CEN6/ARSH4//AmpR//ColE1) | pAG_22-2::(Pscsnr52-22_2-Tsup4//Pagtef1-kan-Tagtef1//2u//CmR//ColE1) | This study |
| FEY_5 | *S.c.* CENPK | - | ChrXV::(Pact1-yEmCitrine-Tadh1//Pagtef1-Nat-Tagtef1) | - | - | - |
| FEY_15 | *S.c.* CENPK | - | HO::(Pgal10-dCas9Mx1-Tspo1//Psptef1-nat-Ttip1) | pY128:(Psnr52-gRNAscr-TtracrSUP4//Pagtef1-kan-Tagtef1//2micron//CmR//ColE1) | - | This study |
| FEY_18 | *S.c.* W303 | ade2-1 ura3-1 his3-11 trp1-1 leu2-3 leu2-112 can1-100 | - | p414-Ptef1-Cas9-Tcyc1 | p426-Psnr52-gRNA_CAN1-Tsup4 | DiCarlo et al., Addgene, Plasmids #43802 & #43803 |
| FEY_27 | *S.c.* S288C | - | hap1:HAP1 | pCSN067::(LbCpf1-SV40//Ptef1-kan-Ttef1//Ptrp1-TRP1-Ttrp1//CEN6/ARSH4//AmpR//ColE1) | - | Verwaal et al., Addgene, Plasmid #101748 |
| FEY_29 | *S.c.* S288C | - | hap1:HAP1 | pUDE722::(Psnr52-Cpf1can1-Tsup4//Ptef1-kan-Ttef1//2u//AmpR//ColE1) | - | Swiat et al., Addgene, Plasmid #101748 |
| FEY_30 | *S.c.* S288C | - | hap1:HAP1 | pSH66::(Pgal1-Cre-Tcyc1//Pagtef1-nat-Tagtef1//CEN6/ARSH4//AmpR//ColE1 | - | Hegemann et al., Euroscarf, P30672 |
| FEY_37 | *S.c.* S288C | - | hap1:HAP1 | pY128:(Ppgk1-pst-Ttdh1//Pagtef1-kan-Tagtef1//2micron//CmR//ColE1) | - | This study |
| FEY_43 | *S.c.* S288C | - | hap1:HAP1 HO::(Psktef1-mdst-Tecm10//Psptef1-nat-Ttip1//Psmtdh3-hfst-Ttdh3//Psmtef1-cast-Teno1) | - | - | - |
| FEY_45 | *S.c.* S288C | - | hap1:HAP1 HO::(Psbtef1-nppst-Tecm10//Psptef1-nat-Ttip1//Psktdh3-lst-Ttdh1) | - | - | - |
| FEY_48 | *S.c.* BY4742 | his3Δ0 leu2Δ0 lys2Δ0 ura3Δ0 | HO::(Psptef1-yEGFP-2A-mRuby-Trps9a//Psptef1-nat-Ttip1) | pCY112::(ccdb//Pagtef1-Klleu2-Tagtef1//CEN6/ARSH4//CmR//ColE1) | - | This study |
| FEY_55 | *S.c.* BY4741 | his3Δ0 leu2Δ0 met15Δ0 ura3Δ0 | - | pKK1112:(Prev1-Venus-Teno2//LEU2//CEN6//KanR-ColE1) | - | This study |
| FEY_74 | Y.l. Po1f | MATA ura3302 leu2270 xpr2322 axp2deltaNU49 XPR2::SUC2 | - | pCRISPRyl(CEN//UAS1B8-TEF(136)-ylCas9-Tcyc1//ColE1//AmpR//sgRNA//Leu2) | | Schwartz et al., Addgene, Plasmid # 70007 |
| FEY_75 | *K.p.* ATCC 76273 | - | - | trp2::(Kan//pUC//attL//Pgap-aMFnoEAEA-RFP-Taox1/Nat/ColE1/Kan//attR) | | This study |

**Table S2.** BLASTN results for all engineered signatures in their respective genome assemblies

*Chromosome integrations*

| Assembly | Percent Identity | # of Gaps | Query Cover |
|---|---|---|---|
| FEY_1 | 99.98 | 1 | 100 |
| FEY_2 | 99.99 | 0 | 100 |
| FEY_5 | 99.76 | 6 | 100 |
| FEY_15 | 99.97 | 1 | 100 |
| FEY_18 | - | - | - |
| FEY_27 | - | - | - |
| FEY_29 | - | - | - |
| FEY_30 | - | - | - |
| FEY_37 | - | - | - |
| FEY_43 | 99.97 | 2 | 100 |
| FEY_45 | 99.91 | 3 | 100 |
| FEY_48 | 99.95 | 1 | 100 |
| FEY_55 | - | - | - |

*Plasmid 1*

| Assembly | Percent Identity | # of Gaps | Query Cover | Length | Correct Length |
|---|---|---|---|---|---|
| FEY_1 | - | - | - | - | - |
| FEY_2 | 98.73 | 37 | 100 | 6143 | 6181 |
| FEY_5 | - | - | - | - | - |
| FEY_15 | 99.8 | 9 | 100 | 4811 | 4820 |
| FEY_18 | 98.46 | 97 | 100 | 10984 | 9524 |
| FEY_27 | 99.93 | 2 | 100 | 11326 | 11322 |
| FEY_29 | 100 | 0 | 100 | 5058 | 5058 |
| FEY_30 | 99.88 | 1 | 100 | 7103 | 7108 |
| FEY_37 | 99.87 | 7 | 100 | 6289 | 6296 |
| FEY_43 | - | - | - | - | - |
| FEY_45 | - | - | - | - | - |
| FEY_48 | 99.98 | 0 | 100 | 4746 | 4746 |
| FEY_55 | 100 | 0 | 100 | 6239 | 6239 |

*Plasmid 2*

| Assembly | Percent Identity | # of Gaps | Query Cover | Length | Correct Length |
|---|---|---|---|---|---|
| FEY_1 | - | - | - | - | - |
| FEY_2 | 99.78 | 5 | 100 | 9654 | 9495 |
| FEY_5 | - | - | - | - | - |
| FEY_15 | - | - | - | - | - |
| FEY_18 | 99.65 | 55 | 100 | 6263 | 6274 |
| FEY_27 | - | - | - | - | - |
| FEY_29 | - | - | - | - | - |
| FEY_30 | - | - | - | - | - |
| FEY_37 | - | - | - | - | - |
| FEY_43 | - | - | - | - | - |
| FEY_45 | - | - | - | - | - |
| FEY_48 | - | - | - | - | - |
| FEY_55 | - | - | - | - | - |

**Table S3.** Evaluation of *de novo* Nanopore assemblers at various genome coverage depths with structure-related metrics

| Coverage | Assembler | N50 Avg (Mb) | # of Contigs Avg | Largest Contig Avg (Mb) |
|---|---|---|---|---|
| 10X | MiniASM | 0.067 | 48.0 | 0.122 |
| | Canu | 0.116 | 126.6 | 0.383 |
| | Flye | 0.220 | 99.4 | 0.752 |
| | SMARTdenovo | 0.074 | 56.0 | 0.141 |
| 20X | MiniASM | 0.247 | 61.6 | 0.789 |
| | Canu | 0.667 | 38.2 | 1.064 |
| | Flye | 0.846 | 25.6 | 1.336 |
| | SMARTdenovo | 0.230 | 64.2 | 0.601 |
| 40X | MiniASM | 0.810 | 19.2 | 1.407 |
| | Canu | 0.858 | 22.4 | 1.411 |
| | Flye | 0.890 | 21.8 | 1.518 |
| | SMARTdenovo | 0.682 | 29.8 | 1.069 |
| 60X | MiniASM | 0.894 | 19.0 | 1.159 |
| | Canu | 0.908 | 22.0 | 1.496 |
| | Flye | 0.901 | 21.0 | 1.525 |
| | SMARTdenovo | 0.745 | 28.0 | 1.500 |
| Reference | | 0.914 | 22.0 | 1.519 |

**Table S4.** Improvement to assembly accuracy using different polishing tools.

| Assembler | Raw | Medaka | Racon | Pilon |
|---|---|---|---|---|
| MiniASM | 86.77 | 99.41 | 99.89 | 99.90 |
| Canu | 99.33 | 99.42 | 99.90 | 99.91 |
| Flye | 98.16 | 99.41 | 99.90 | 99.90 |
| SMARTdenovo | 99.13 | 99.39 | 99.89 | 99.89 |

**Table S5.** Improvement to assembly accuracy through polishing at various genome coverage depths of Illumina reads.

| Coverage | Assembler | Avg ID | # of SNPs |
|---|---|---|---|
| 0X | MiniASM | 99.41 | 3629 |
| | Canu | 99.42 | 3714 |
| | Flye | 99.41 | 3639 |
| | SMARTdenovo | 99.39 | 3589 |
| 5X | MiniASM | 99.56 | 16824 |
| | Canu | 99.56 | 16879 |
| | Flye | 99.56 | 16915 |
| | SMARTdenovo | 99.54 | 16749 |
| 10X | MiniASM | 99.68 | 14149 |
| | Canu | 99.69 | 14141 |
| | Flye | 99.69 | 14202 |
| | SMARTdenovo | 99.68 | 13848 |
| 20X | MiniASM | 99.82 | 6361 |
| | Canu | 99.83 | 6281 |
| | Flye | 99.83 | 6340 |
| | SMARTdenovo | 99.80 | 6277 |
| 40X | MiniASM | 99.89 | 1944 |
| | Canu | 99.89 | 1837 |
| | Flye | 99.89 | 1871 |
| | SMARTdenovo | 99.87 | 1813 |
| 100X | MiniASM | 99.89 | 1137 |
| | Canu | 99.90 | 1036 |
| | Flye | 99.90 | 1139 |
| | SMARTdenovo | 99.89 | 1079 |
| 200X | MiniASM | 99.90 | 1027 |
| | Canu | 99.91 | 1027 |
| | Flye | 99.90 | 1081 |
| | SMARTdenovo | 99.89 | 1046 |
| 267X | MiniASM | 99.90 | 1067 |
| | Canu | 99.91 | 981 |
| | Flye | 99.90 | 1049 |
| | SMARTdenovo | 99.89 | 998 |
| Reference | | 99.90 | 785 |

**Table S6.** BUSCO genome completeness assessment for each Nanopore *de novo* assembler at each genome coverage depth and polishing software.

| Coverage | Assembler | Raw | Medaka | Racon | Pilon |
|---|---|---|---|---|---|
| 10X | MiniASM | 0.00 | 2.40 | 17.07 | 17.36 |
| | Canu | 0.00 | 11.28 | 91.00 | 93.69 |
| | Flye | 0.06 | 13.09 | 95.73 | 98.13 |
| | SMARTdenovo | 0.00 | 10.11 | 94.51 | 97.37 |
| 20X | MiniASM | 1.81 | 25.37 | 71.30 | 71.65 |
| | Canu | 8.53 | 40.62 | 96.73 | 97.43 |
| | Flye | 19.23 | 43.54 | 98.01 | 98.36 |
| | SMARTdenovo | 23.44 | 33.49 | 97.43 | 98.25 |
| 40X | MiniASM | 1.11 | 21.45 | 94.56 | 95.50 |
| | Canu | 2.75 | 22.79 | 97.84 | 98.54 |
| | Flye | 3.80 | 18.29 | 97.78 | 98.19 |
| | SMARTdenovo | 4.15 | 19.11 | 97.72 | 98.36 |
| 60X | MiniASM | 0.29 | 6.66 | 27.94 | 28.17 |
| | Canu | 4.73 | 30.98 | 91.93 | 92.46 |
| | Flye | 15.37 | 38.92 | 98.07 | 98.19 |
| | SMARTdenovo | 24.90 | 39.98 | 97.78 | 98.31 |
| Reference | | 98.25 | | | |

**Table S7.** Percentage of *S. cerevisiae* S288C CDS found in each Nanopore *de novo* assembler at each genome coverage depth and polishing software.

| Coverage | Assembler | Raw | Medaka | Racon | Pilon |
|---|---|---|---|---|---|
| 10X | MiniASM | 18.99 | 20.41 | 20.64 | 20.56 |
| | Canu | 6.42 | 95.07 | 95.67 | 95.73 |
| | Flye | 6.55 | 99.18 | 99.49 | 99.47 |
| | SMARTdenovo | 6.18 | 98.28 | 98.86 | 98.98 |
| 20X | MiniASM | 11.02 | 75.23 | 75.60 | 75.43 |
| | Canu | 16.77 | 99.13 | 99.18 | 99.11 |
| | Flye | 20.44 | 99.40 | 99.39 | 99.39 |
| | SMARTdenovo | 99.59 | 99.69 | 99.76 | 99.69 |
| 40X | MiniASM | 80.77 | 97.46 | 97.65 | 97.71 |
| | Canu | 98.43 | 99.27 | 99.40 | 99.40 |
| | Flye | 98.79 | 99.27 | 99.44 | 99.51 |
| | SMARTdenovo | 98.79 | 99.20 | 99.35 | 99.39 |
| 60X | MiniASM | 22.95 | 31.26 | 31.39 | 31.38 |
| | Canu | 87.51 | 94.22 | 94.35 | 94.30 |
| | Flye | 98.60 | 99.13 | 99.16 | 99.18 |
| | SMARTdenovo | 98.89 | 99.40 | 99.45 | 99.47 |
| Reference | | 99.66 | | | |

#### Supplementary Figures

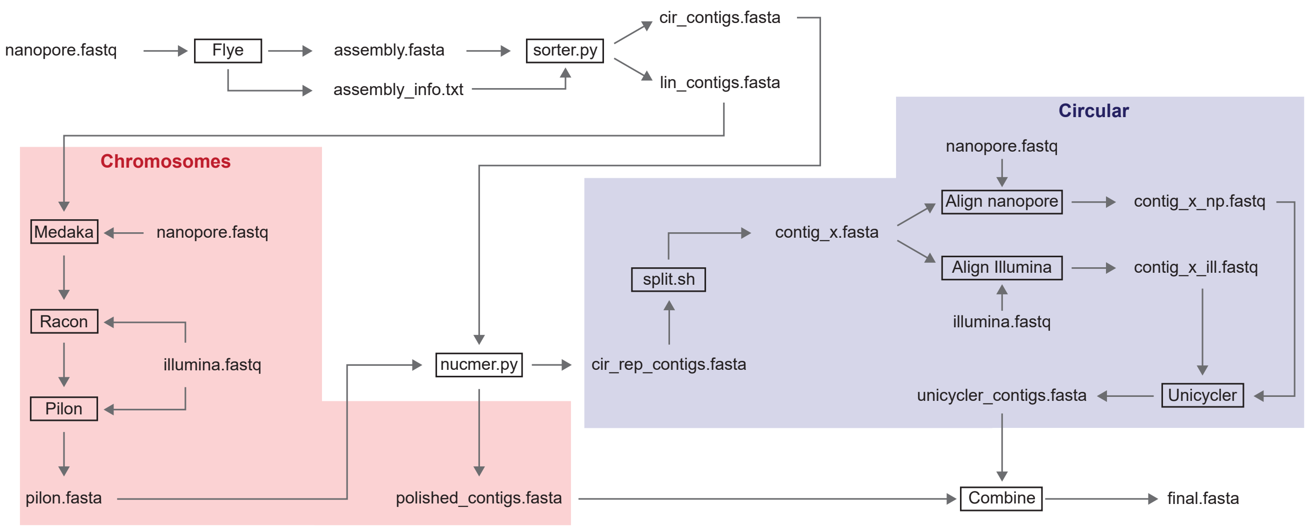


**Figure S1.** Detailed PRYMETIME genome assembly pipeline. Yeast chromosomes and plasmids were assembled and polished in two separate steps: chromosomes were assembled using the Flye assembler followed by polishing with Medaka, Racon, and Pilon, while plasmids and circular elements were extracted and re-assembled using Unicycler. Chromosomes and plasmids were then combined to yield the final assembly file.

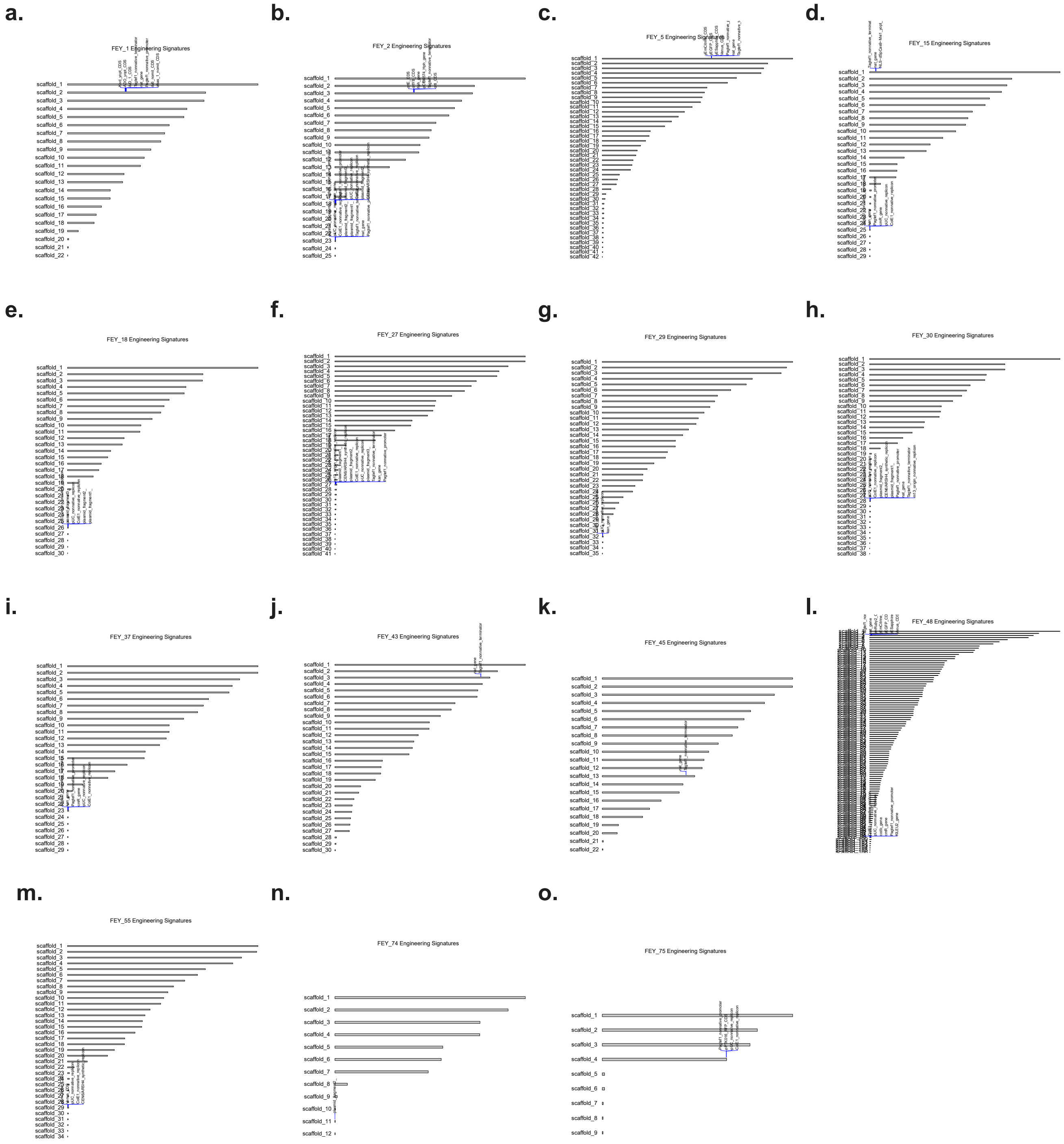


**Figure S2.** Genome plots of each engineered strain and their associated non-native engineering signatures. **a.** *S. cerevisiae* FEY_1. **b.** *S. cerevisiae* FEY_2. **c.** *S. cerevisiae* FEY_5. **d.** *S. cerevisiae* FEY_15. **e.** *S. cerevisiae* FEY_18. **f.** *S. cerevisiae* FEY_27. **g.** *S. cerevisiae* FEY_29. **h.** *S. cerevisiae* FEY_30. **i.** *S. cerevisiae* FEY_37. **j.** *S. cerevisiae* FEY_43. **k.** *S. cerevisiae* FEY_45. **l.** *S. cerevisiae* FEY_48. **m.** *S. cerevisiae* FEY_55. **n.** *Y. lipolytica* FEY_74. **o.** *K. phaffii* FEY_75.

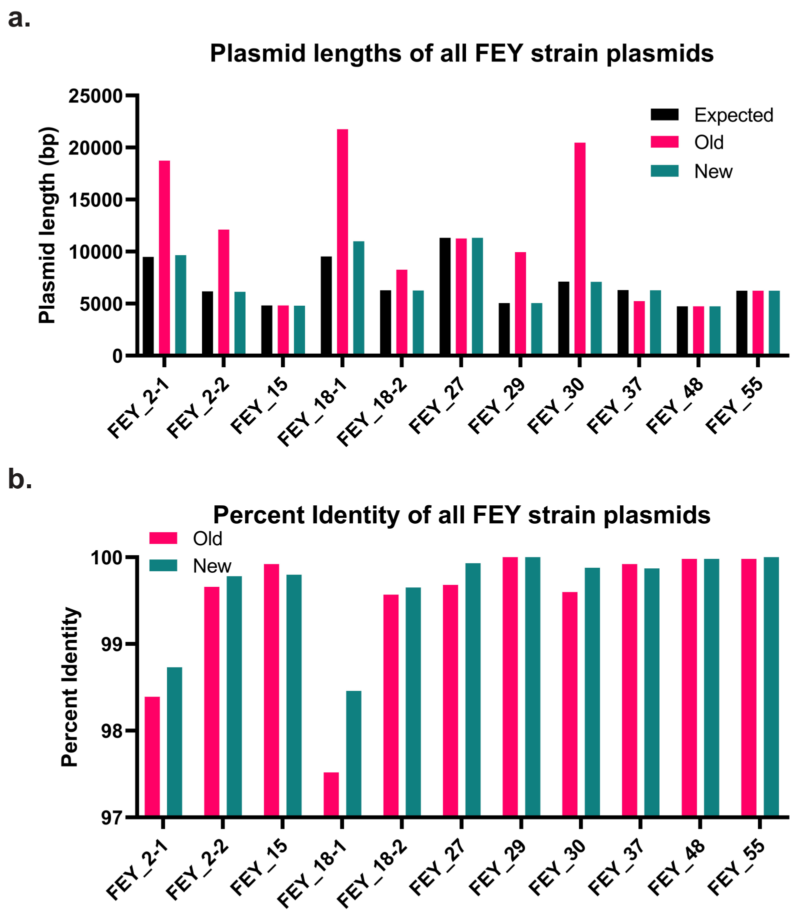


**Figure S3.** Comparison of engineered strain assembly plasmids with or without the added Unicycler. **a.** Length of each engineered plasmid from each genome assembly strategy against the expected length. **b.** BLASTN percent identity of each engineered plasmid from each genome assembly strategy.

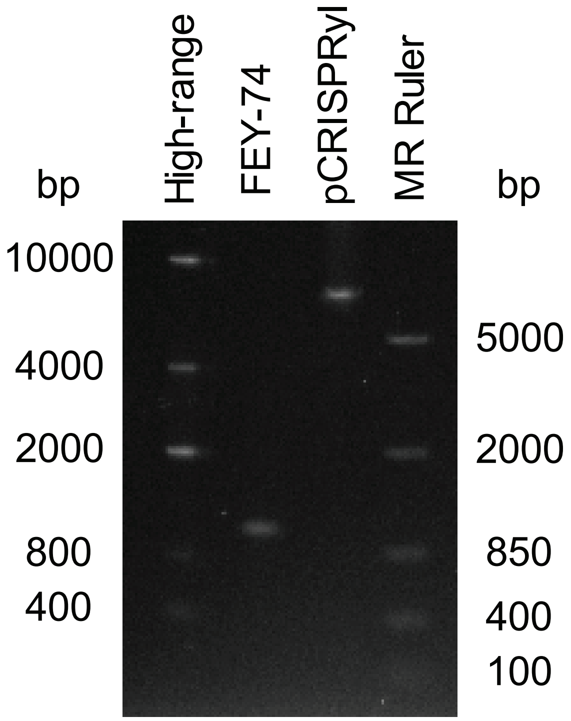


**Figure S4.** A DNA gel showing a PCR across the Cas9 transcription unit in the FEY_74 strain versus the original pCRISPRyl plasmid.

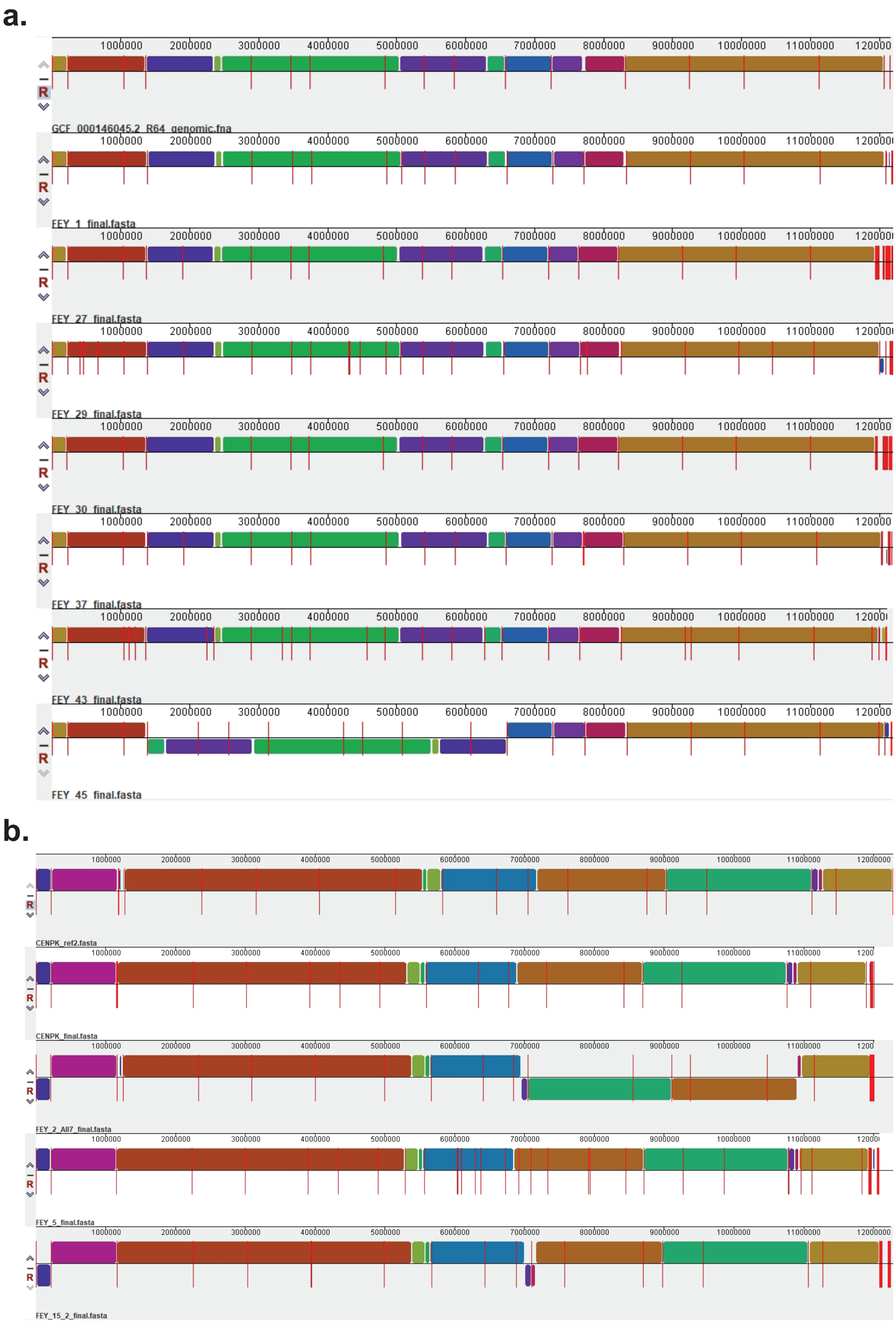


**Figure S5.** Whole genome comparison of engineered *S. cerevisiae* strains against their respective parent strains using Mauve. **a.** Comparison of FEY_1, FEY_27, FEY_29, FEY_30, FEY_37, FEY_43, and FEY_45 against a reference S288C assembly. **b.** Comparison of FEY_2, FEY_5, and FEY_15 against two CEN.PK113-7D reference assemblies, including one from this study.

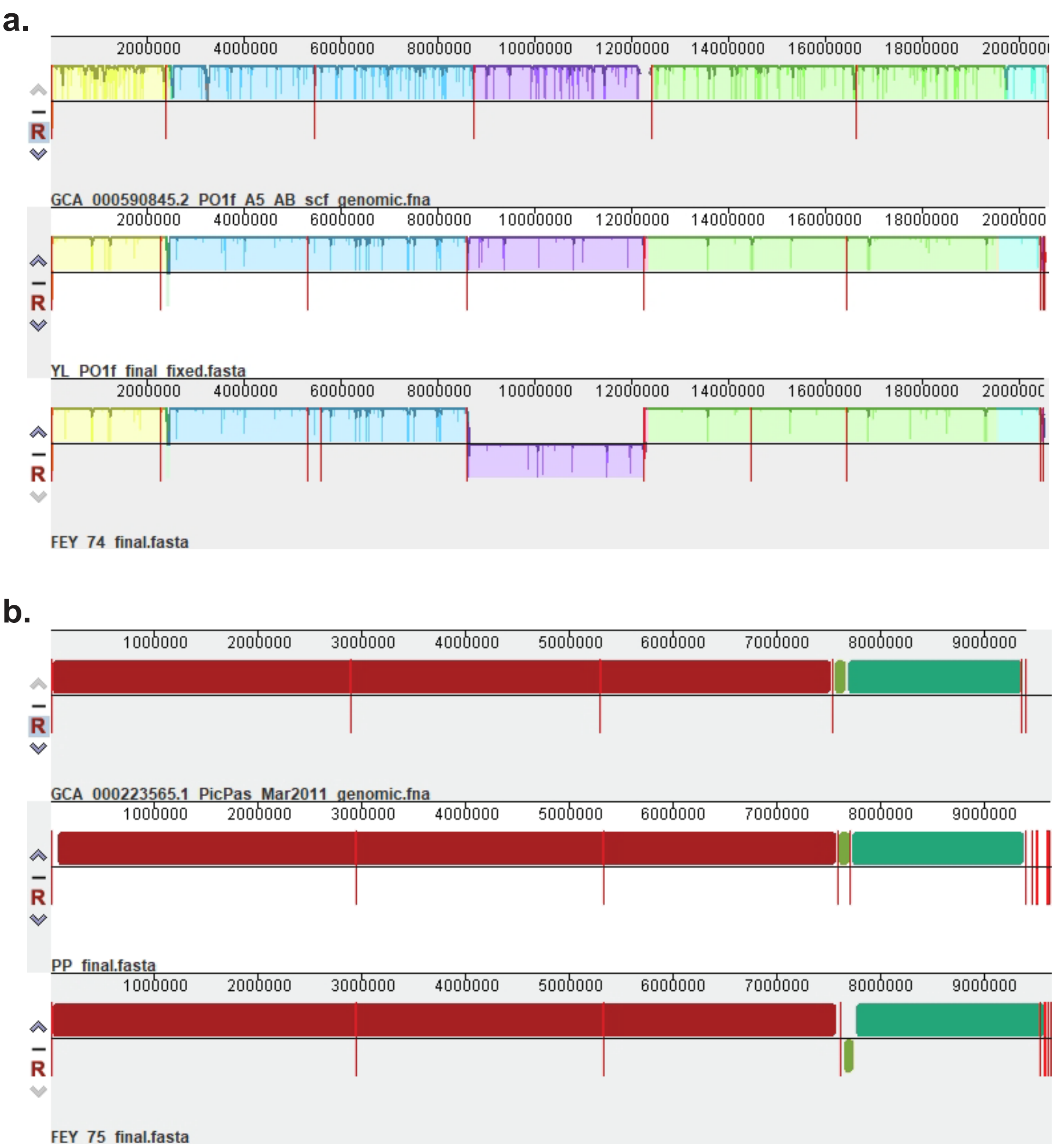


**Figure S6.** Whole genome comparison of engineered nonconventional strains against their respective parent strains using Mauve. **a.** Comparison of FEY_74 against two *Y. lipolytica* PO1f reference assemblies, including one from this study. **b.** Comparison of FEY_75 against two *K. phaffii* CBS 7435 reference assemblies, including one from this study.
